# Quantifying the organization and dynamics of *M. smegmatis* morphology from Long-Term Time-Lapse Atomic Force Microscopy

**DOI:** 10.1101/2025.01.17.633629

**Authors:** Clément Soubrier, Anotida Madzvamuse, Haig Alexander Eskandarian, Khanh Dao Duc

**Affiliations:** Department of Mathematics, University of British Columbia, Vancouver, BC V6T 1Z4, Canada; Blavatnik Institute, Department of Systems Biology, Harvard Medical School, Boston, MA 02115, USA

## Abstract

The rod shaped *Mycobacterium smegmatis* displays complex cell surface morphology, characterized by wave-form cell surface features and driven by asymmetric growth dynamics. To systematically analyze these morphological variations, we developed a comprehensive computational pipeline for automated processing of Long-Term Time-Lapse Atomic Force Microscopy (LTTL-AFM) images of *M. smegmatis* cells cultured in axenic conditions of growth and stress. Upon running the pipeline to produce large enough datasets of single cell height profiles, we identify and statistically study key features that govern cell surface morphology: We confirm that *M. smegmatis* cells undergo bi-phasic, asymmetric pole growth with constant elongation rate at the old pole and a shift in the rate of elongation after a lag phase at the new pole. Stable wave-form cell surface peaks and troughs propagate along the long axis of the cell, which emerge as a result of polar elongation. Backtracking in time from cell division, we detect that division-site selection occurs at the wave-trough nearest mid-cell. To reproduce the fundamental cell features observed, we introduced a reaction-diffusion mathematical model on an evolving one-dimensional surface. Our simulations indicate that the dynamic manifestation of wave-form cell surface morphology in *M. smegmatis* can be explained by the interaction of as few as two “chemical species”, providing a plausible theoretical basis for how molecular determinants may functionally control wave-form morphology in pole-growing bacteria.

## 1 Introduction

*Mycobacterium smegmatis* is a rapidly growing, non-pathogenic bacterium, which is used as a biological model organism for studying the cell physiology of pathogenic mycobacteria [1], such as *M. tuberculosis, M. abscessus* or *M. leprae*, for which genetic manipulation and molecular mechanisms remain poorly understood. While its cell surface morphology has traditionally been characterized as rod-shaped, recent studies revealed a wave-form cell surface morphology and complex pole elongation dynamics, culminating in morphological heterogeneity at the individual cell level within an isogenic clonal population [2, 3]. As the amplitude of the wave-form patterns (∼ 100 *nm*) requires high resolution imaging beyond the optical diffraction limit, detecting such morphological variations required the development and use of Long-Term Time-Lapse Atomic Force Microscopy (LTTLAFM) [4]. This technique represents a multi-parametric scanning probe imaging modality enabling discovery-based studies of bacterial cell surface physiology resulting in hypothesis-generating characterizations of fundamental cell processes that are otherwise immeasurable [4, 5, 6]. For mycobacterial cell physiology, expanding the breadth of phenomenological understanding provides a new paradigm for potentially identifying how molecular factors are relevant in influencing fundamental cell processes.

LTTL-AFM studies have provided novel insights into a variety of fundamental mycobacterial cell processes for which molecular mechanisms remain unknown. LTTL-AFM imaging has enabled an expanded understanding for how proliferative cell processes take place, such as growth, division, and adaptation [2, 3, 7]. However, the nature of these discoveries is constrained by the analysis of cell features extracted from manually segmented cells, resulting in limited throughput and systematization of data generation. Computational and mathematical analyses were prohibited by manual data processing and analysis. While existing bioinformatic tools (e.g. [8, 9, 10]) can be combined to identify, segment and extract biophysical properties of single cells from large images, these methods typically require fine tuning and parameter-calibration. These steps are further confounded by the different nature of LTTL-AFM images. Specific artifacts and microscope parameters sporadically emerge over the duration of an experiment. As a result, previous characterizations of *M. smegmatis* cell properties have predominantly relied on manually curated LTTL-AFM datasets [3] potentially introducing inherent subjective bias and limited sample sizes.

In the present study, we develop a computational pipeline for processing LTTL-AFM datasets, by automating cell segmentation, tracking of cells and division events, and outputting key geometrical features over time. As we ran the pipeline to collect large sample of single cell images, we further conducted various statistical analyses of *M. smegmatis* morphological features, their dynamic emergence and phenotypic heterogeneity at cellular and sub-cellular levels. We validate the consistent presence of bi-phasic and asymmetric pole elongation dynamics (previously observed in [2]), and study the morphology and position of the cell division site. These systematic studies of cell shape dynamics finally led us to formulate a mathematical reaction-diffusion model that can produce and maintain similar wave-form patterns over time, suggesting how molecular determinants can control mycobacterial cell processes of growth and division.

## 2 Material and Methods

Protocols for data acquisition and processing are briefly summarized in the following section, with more details provided in SI file (Section S1)

### 2.1 LTTL-AFM raw data

*M. smegmatis* bacilli were non-specifically immobilized on thinly spin-coated PDMS coverslips as per [3]. *M. smegmatis* was cultured in 7H9 growth medium at 37°C. Perfusion system was used for fluidic exchange, temperature control, and air bubble trap. Antibiotic treatments were conducted by supplementing antibiotics in 7H9 growth medium. The raw AFM data was recorded using a Bruker AFM, with a period of 9 minutes per image or more, and covering an area of roughly 100 - 1000 *μm*^2^. Physical channels recorded include height sensor, peak force error and Derjaguin–Muller–Toporov (DMT) modulus, representing the stiffness. For more detail on the data acquisition see [3, 11]. During pre-processing, we extracted the AFM log files and erased artifacts using the pySPM library, before flattening and re-dimensioning the images at the same size.

### 2.2 Cell segmentation and tracking

We ran CellPose [8] to detect and segment individual cells in each image. Multiple physical data images contained in log files (accounting for stiffness, peak force error and height) were combined to improve segmentation. Upon obtaining time-series data of cell masks, single cell tracking and division events detection were performed using a custom algorithm, that builds a graph from a pairwise geometric score function measuring cell similarity. Longest paths inside each connected component were extracted to produce a lineage tree (see SI file Section S1.2 for more details).

### 2.3 Characterization and measurement of morphological features

Cell centerlines were derived from the skeleton of the associated masks, after pruning and extending the skeleton to the cell poles. The height profile of each cell was then obtained by measuring the height along the centerline. Similarly stiffness height profiles were also measured and averaged over sub-polar and central cell regions. Height profile features, namely peaks and troughs, were detected as local maxima and minima of the smoothed height profile, respectively. Using lineage information, we produced kymographs by aligning height profiles and tracked peaks and troughs along the centerline over time and across generation (for more details, see SI file Sections S1.3 and S1.4). This alignment also enables to locate division sites and track their properties back in time.

We measured the cell growth using time series of aligned centerline lengths. The “New-End Take-Off”(NETO) [2, 12] was then inferred by fitting a piecewise affine function using least square regression. Once the NETO was computed, we fitted the elongation growth curve at each pole (old and new) with piecewise affine functions and used the associated slope coefficients to measure elongation speed pre- and post-NETO. After division, we aligned daughter kymographs to the mother kymographs, and computed the division site by averaging the new pole coordinates of daughter cells. We computed the local curvature over a window of 0.4 *μm* around the division site, which represents approximately half the size of a peak or trough.

### 2.4 Mathematical modeling of height profile dynamics

We model the observed height profile dynamics using a coupled system of partial differential equations, defined over an interval Ω(*t*) = [−*L*_1_(*t*); *L*_2_(*t*)] ⊂ ℝ evolving in time according to the bi-phasic asymmetrical growth observed in the data. We work on a fixed time interval *I* = (0, *T*) with *T* the final simulation time. Here, *L*_1_(*t*) represents the old pole position and *L*_2_(*t*) the new pole, with initial conditions *L*_1_(0) = 0 and *L*_2_(0) = 1. The pole growth is piecewise constant so that, for posi-tive time 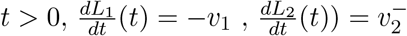 before NETO and 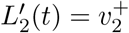 after NETO. The polar regions are assumed to have constant length *δ*, typically 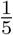 to 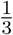 of the cell length. The system of PDE’s follows an activator-depleted model, also known as the Schnakenberg model [13], describing non linear spatio-temporal interactions between two chemical species, with concentrations *u*(*x, t*) and *v*(*x, t*) (*x* ∈ Ω(*t*) and *t* ∈ *I*). The system of PDE’s on the one-dimensional evolving domain Ω(*t*) reads:

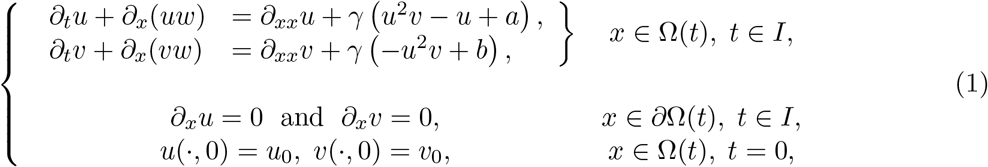

where *a* and *b* are two constants accounting for the production rates of *u* and *v* respectively, *d* is the ratio of the two species diffusivity constants and *γ* represents the interaction intensity. If not otherwise stated, we used parameters *a* = 0.1, *b* = 0.9, *γ* = 800 and *d* = 10 in the figures. The scalar field *w*(*x, t*) is the material velocity and captures the growth of the cell. We chose *w*(*x, t*) to reproduce the observed polar cell elongation dynamics, that is null over the central region of the cell *x* ∈ [−*L*_1_(*t*) + *δ*; *L*_2_ − *δ*] and linear over the polar regions:

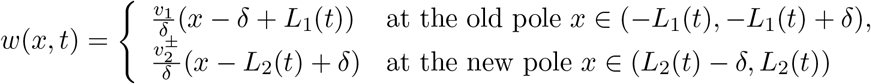

We numerically solved this PDE [14] using an evolving surface finite element method as described in SI file Section S2, and implemented with the Python package FEniCSx (based on [15, 16, 17]). Both fully implicit and implicit-explicit (IMEX) numerical schemes were implemented and compared to ensure convergence, following both mesh and time-step refinements.

### 2.5 Datasets

Our dataset contains four subsets of wild type mc2 55 *M. smegmatis* strands in various conditions. Among these datasets, one describes bacteria that are separately exposed to the antibiotics INH.

Overall, there are 502 frames, with 12 566 detected masks, and 336 tracked cells. Upon performing each statistical analysis, we filtered the data to keep high quality samples (see SI file Section S4). The original dataset and the processed datasets are both available at this OSF (link).

### 2.6 Implementation and code availability

The pipeline for the automated treatment of AFM data and statistical analysis, were implemented in Python 3.10. Our code and scripts are available to access on Github (link) and on OSF (link).

## 3 Results

### 3.1 Automated pipeline for extracting morphological trajectories and features of single *M. smegmatis* cells in LTTL-AFM

To statistically study the morphological properties and dynamics of *M. smegmatis*, we processed and analyzed large datasets of LTTL-AFM data using the pipeline described in Section 2, and schematized in Figure 1. This pipeline corrects for artifacts due to aberrations in scanning probe measurements, manifesting in AFM height images as scars and saturation, (Figure 1A) in which the dotted dashed box representing a scar that is detected and removed. The pipeline automatically selects individual cell shapes with high quality (Figure 1A), to output cell outlines, centerlines and centroids as key features to be analyzed downstream. Automated tracking of cells over time and over divisions produces lineage trees, which subsequently enable to study the dynamics of cell features during a full cell cycle and across generations (Figure 1B). We constructed height profiles from the centerline that represent the diameter of the cell along its longitudinal axis. The tracking of time series data yields a kymograph of height profiles for each individual cell, which are aligned by optimizing congruity of wave-form peaks and troughs (Figure 1B and S4). We report how our pipeline is used to study various features of *M. smegmatis*, namely its asymmetric and bi-phasic polar growth, the presence of peaks and troughs and location of the division site.

**Figure 1.**
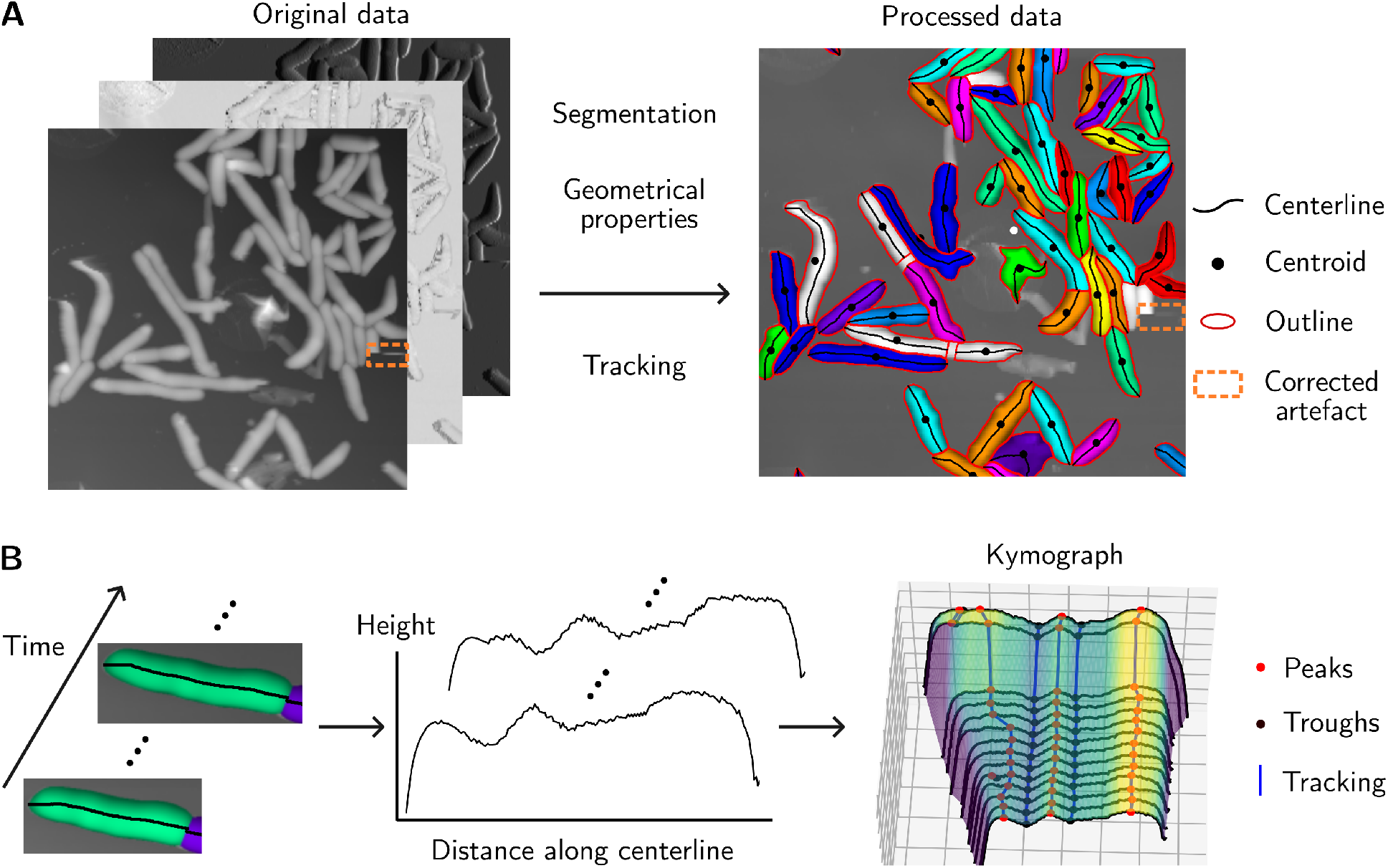
Schematic representation of the pipeline, from microscope data to aligned kymographs. (A) Cell segmentation and tracking from the raw data. The original AFM data is composed of several physical channel. In the yellow dashed box, a scar is detected and removed. Grey masks are ignored because of bad morphology or insufficient tracking data. (B) For each generation, centerlines are computed over the cell life. Height profiles are then extracted and aligned. Finally, peaks and troughs are detected, tracked and kymographs are produced.

### 3.2 Detection of bi-phasic and asymmetric cell growth

*M. smegmatis* growth dynamics is characterized by asymmetric polar elongation. While the old pole (pre-existent from the previous generation) exhibits a constant elongation velocity, the new pole (formed by the previous division) exhibits a bi-phasic growth dynamic characterized by slow growth, followed by a shift to rapid growth converging on symmetric polar elongation [2]. The bi-phasic growth dynamic is reminiscent of pole growth in the fission yeast, *Schizosaccharomyces pombe*, and previously characterized as “New-End Take-Off” (NETO) [12]. Kymograph and lineage information was used to statistically quantify bi-phasic growth dynamics in *M. smegmatis* (Figure 2A). For every cell with complete lineage information, we computed the overall (black), old pole (red) and new pole elongation (blue) velocities, as well as when NETO occurs (Figure 2B, S1A).

**Figure 2.**
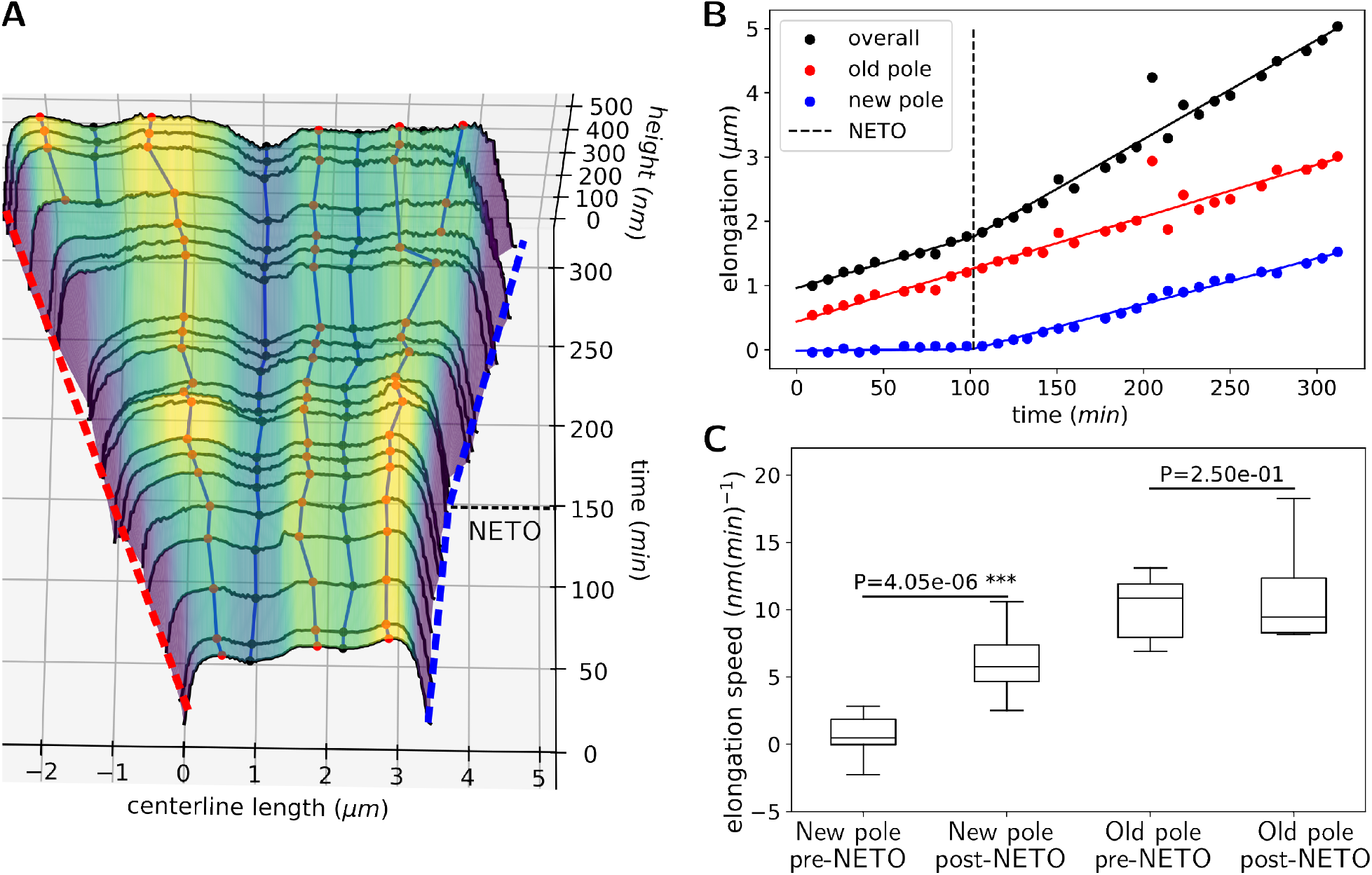
Detection and quantification of the bi-phasic pole growth dynamic. (A) Kymograph of processed and aligned height profiles, from cell birth to division. Polar elongations (blue and red dashed lines) are extracted from pole locations. The new pole (blue) has a biphasic growth with a “New-End-Take-Off” (NETO) at ∼ 150 minutes. (B) Example of piecewise linear fit of overall cell elongation (black) and extraction of NETO time, with old pole (red) and new pole (blue) dynamics. (C) Bar plots of elongation speed for old and new poles over pre- and post-NETO times.

Similar distributions of old pole elongation velocities are observed before and after NETO whereas elongation velocity is slow during the lag phase (pre-NETO) at the new pole and increases to a velocity approaching that observed at the old pole, post-NETO (Figure 2C). Convergence of elongation velocities at both old and new poles is maintained long after NETO (suggested by Figure 2B and S1C, with a non significant p-value of 0.082 for *n* = 52 cells).

We asked how antibiotic perturbation driving growth arrest impacts the bi-phasic growth dynamic. Isoniazid (INH) represents a bactericidal, first-line antibiotic for treating *M. tuberculosis* infection. INH treatment results in reduced mycolate biosynthesis, a critical component of the outer cell wall [18]. INH generally provokes growth arrest as well as reductive division, and swelling [3]. We observed that growth arrest is characterized by an equal reduction in elongation velocity of both the old and new poles, to pre-NETO growth levels (Figure S1B). NETO was abolished at the new pole upon INH treatment, culminating in a persistent lag phase, which is characterized by a slow elongation velocity (Figure S1B). Pole growth asymmetry requires a lag phase, pre-NETO, wherein new and old poles elongate at different rates. Control of pole elongation asymmetry represents a potential driver of phenotypic heterogeneity in an isogenic population.

### 3.3 Wave-form morphology is established in sub-polar zones of elongating poles

Previous investigations using LTTL-AFM revealed a wave-form cell surface morphology propagating in a semi-regular manner along the long-axis of *M. smegmatis* cells [3]. Wave-peaks and wave-troughs represent consistent variations of the cell diameter and are inherited over generations. We extracted waveform features from our datasets and performed statistical analyses to quantify aspects of their emergence and propagation along the length of the cell surface. The average distance between two wave-troughs is ∼ 1.5*μm* and the average amplitude between a wave-trough and wave-peak is ∼ 100*nm* (Table 1, Figure S2A). The wave-form amplitude represents nearly 10% − 20% of the diameter of the cell along its short axis, suggesting that the volume harboring intra-cytosolic components likely favors wave-peaks. Wave-form patterns are created with semi-regularity over the course of the cell cycle. The number of wave-peaks increases from on average 2.6 at birth to 3.9 at cell cleavage; the number of wave-troughs is lower, on average 1.6 at birth and 2.9 at cell cleavage (Table 1). Since division usually takes place at a wave-trough and cell poles count as wave-peaks, cells exhibit one fewer wave-trough.

**Table 1:** Statistics of *M. smegmatis* cell surface wave-patterns, including their number, amplitude, distance and creation location. The creation location represents the distance to the closest pole, relative to the cell length. Wave-peaks and wave-troughs are created at sub-polar regions.

|  | Number<br>at birth | Number<br>at division | Number<br>created<br>during life | Amplitude<br>(nm) | Distance<br>(μm) | Creation<br>location<br>(% cell length) |
| --- | --- | --- | --- | --- | --- | --- |
| wave-peaks | $2.6 \pm 0.7$ | $3.9 \pm 1.2$ | $1.4 \pm 1.3$ | $75 \pm 93$ | $1.60 \pm 0.67$ | $14 \pm 7$ |
| wave-troughs | $1.6 \pm 0.7$ | $2.9 \pm 1.2$ | $1.3 \pm 1.2$ | $69 \pm 63$ | $1.48 \pm 0.54$ | $17 \pm 6$ |

Spatiotemporal tracking of mycobacterial cell surface topography enabled us to identify when and where new wave-form features are created (Figure 3A). Establishment of new morphological features occurs within a range of space comprising 15% of the cell length emanating inwards from the poles (Table 1). Elongation-driven cell surface expansion outward sets the creation of new wave-form troughs and peaks within a 1.5*μm* space near each pole. We computed the variations in feature height and position along the cell length over time (Figure 3B). In the central cell region, variation in height is small suggesting that morphological features are stable along the cell cycle. The sub-polar space is characterized by increased morphological variation (Figure 3B, S2C) and decreased mean cell surface stiffness (Figure S2D), as compared to mid-cell (Figure 3C).

**Figure 3.**
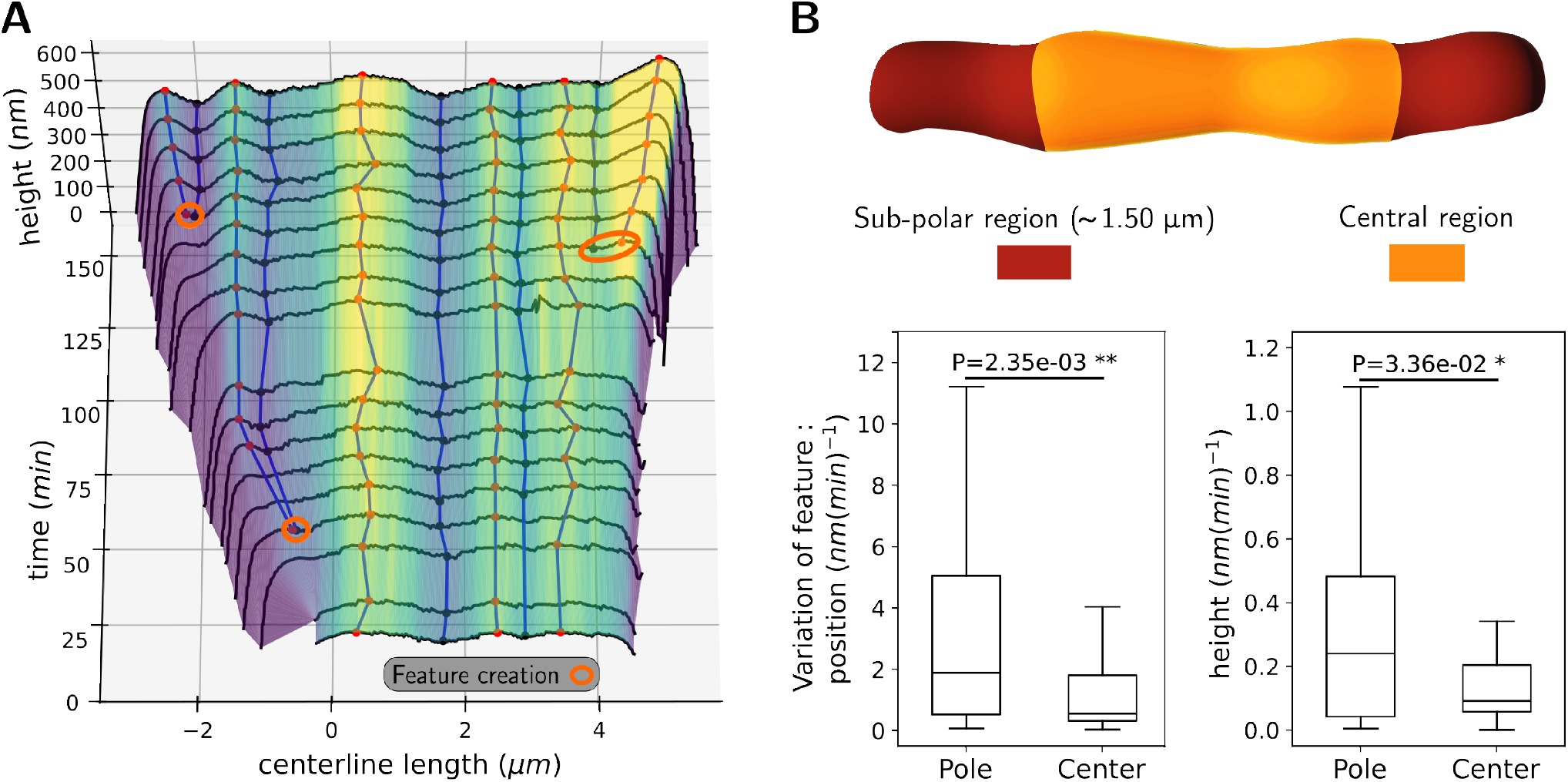
Characterization of two distinct morphological regions in *M. smegmatis*: the central and sub-polar region. (A) Morphological features are extracted from cell kymographs, such as the spatial localization and amplitude of wave-peaks and wave-troughs, or their creation (orange). (B) Box-plot comparing variations of pattern properties (position and height) in the sub-polar and central regions (cartoon bacteria adapted from [7]). Morphological variations are higher at the poles.

We asked whether pole growth asymmetry influences cell surface morphology. Variation in inter-feature amplitudes and distances are similar for both old and new poles (Figure S2C). However, reduced new pole elongation during the lag phase typically results in the addition of only 100 *nm*, which is insufficient to harbor the creation of a wave-form feature: over 32 detected pattern creation, none happened at the new pole before NETO. While new morphological features are not created at the new pole during the lag phase, pre-existent features are mechanically reinforced and morphologically stabilized.

### 3.4 Characterization and timing of division site selection

What fundamental principles govern division site selection in *M. smegmatis*? While molecular determinants of division site selection in *M. smegmatis* remain largely unknown, LTTL-AFM imaging previously revealed that division site selection occurs at the wave-trough nearest mid-cell [3]. To expand our understanding of this process, we aimed to locate and study the properties of the division site from the kymographs. Using lineage tree information, we aligned daughter cell kymographs with their mothers, marked the division site location (yellow pentagon in Figure 4A) and traced its position back in time until cell birth (yellow star). Table 2 summarizes key features of the division site, including its distance to the closest wave-form feature, amplitude, and relative position.

**Table 2:** Division site geometrical properties and prediction for its selection time. The p-values corresponds to a 2 sample and 1 sample T-tests. For the division position, the null hypothesis assumes a mean at the cell center (50%), with 0% representing the new pole.

| | Distance to closest peak ($\mu\text{m}$) | Distance to closest trough ($\mu\text{m}$) | Closest feature amplitude (nm) | Overall feature amplitude (nm) | Division position (% cell) | Predicted selection time $T_0$ (% life) |
| --- | --- | --- | --- | --- | --- | --- |
| mean ± std | $0.95 \pm 0.55$ | $0.71 \pm 0.69$ | $112 \pm 93$ | $82 \pm 81$ | $44 \pm 5$ | $36 \pm 29$ |
| p-value test | $0.039*$<br>Student | | $3.8 \cdot 10^{-5}***$<br>Student | | $1.4 \cdot 10^{-3}**$<br>Student | |

**Figure 4.**
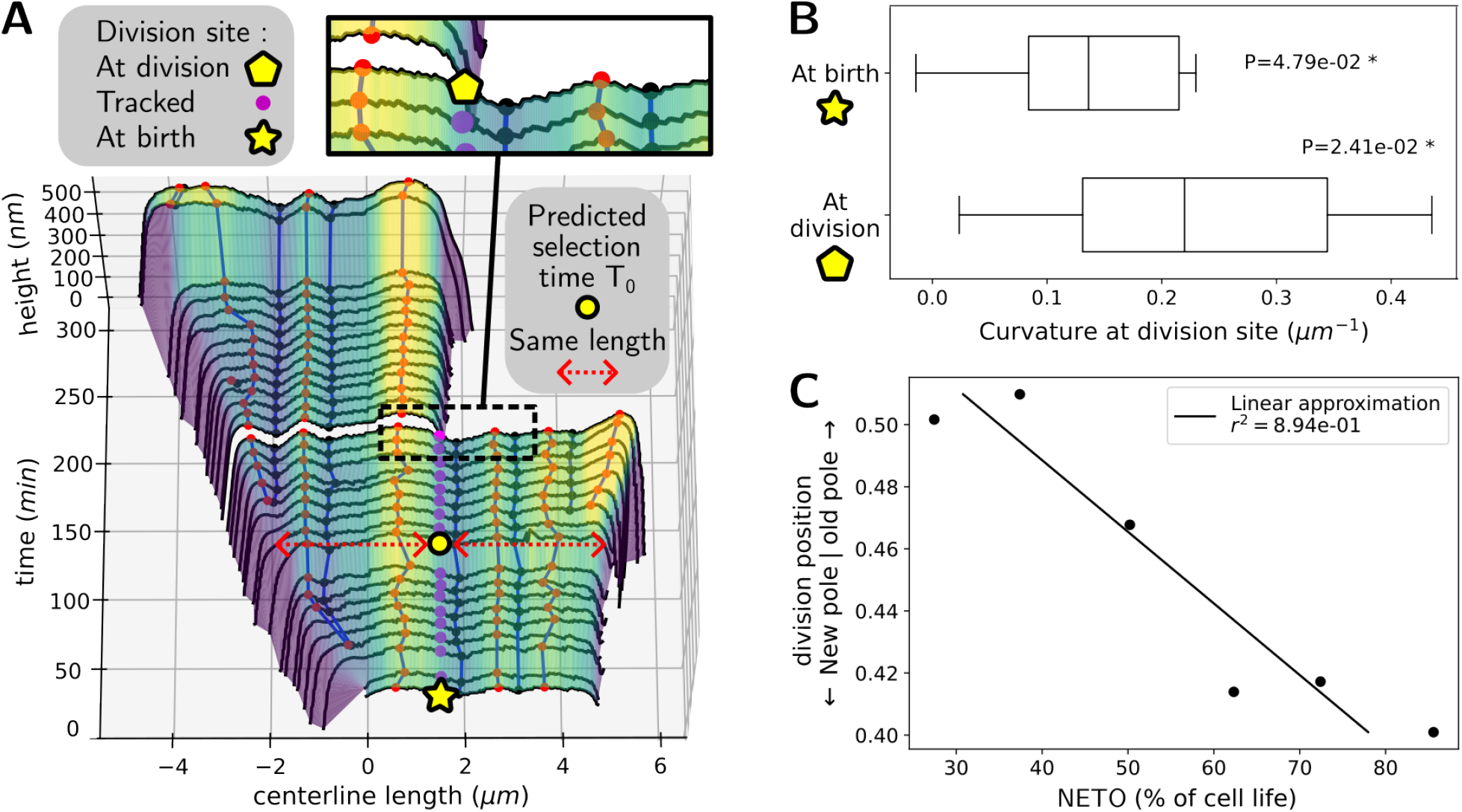
Quantification of division site morphology and selection timing. (A) Kymographs of mother and daughter cells representing the division site tracked back in time. We predict a division selection time *T*_0_ as the moment at which the back-tracked division site is located in the middle of the cell. (B) Representation of the division site curvature at division, and when tracked back to birth. By convention, troughs have positive curvature. The plotted p-value corresponds to a one sample T-test, with zero curvature mean null hypothesis. (C) Plot of relative division position, as a function of the timing of NETO.

Our analyses suggest that division sites consistently form closer to wave-troughs than wave-peaks (0.71 *μm*, compared to 0.95 *μm* on average, with a p-value under 5%). The height differential between wave-peaks and wave-troughs surrounding division sites was significantly larger compared to other regions of the cell (112 *μm*, compared to 82 *μm* on average, with a p-value under 0.1%), with surrounding wave-form features having similar amplitude (see Figure S3B). The division site height profile exhibits positive local curvature (from in intra-cytosolic vantage) (Figure 4B) and slightly increases following the formation of the pre-cleavage furrow and prior to cell cleavage of cytokinesed sibling cells [11]. We detected a spatial bias for division site selection towards the new pole, as it is located on average at 44 ± 5% of the cell length from the new pole (Table 2). We confirmed this bias, by performing a 1 sample T-test rejecting the null hypothesis that division occurs near mid-cell, with a p-value of 0.004. In comparison to untreated wildtype cells, symmetrically elongating INH-treated cells (for which NETO is abrogated) (Figure S1B) harbor division sites located closer to mid-cell, at 50 ± 6% of the cell length (n = 7) (Figure S3A), with the same T-test now yielding a p-value of 0.88.

What is the relationship between the division site and bi-phasic pole growth dynamics in *M. smegmatis*? To investigate the relationship between NETO timing and division site selection, we performed a linear fit of NETO timing to the spatial position of the division site. A longer lag phase (delayed NETO) correlates with the division site selection occurring closer to the new pole (with a coefficient of determination *r*^2^ of 0.89, and a slope of −2.3 · 10^−3^) (Figure 4C). A short lag phase, at ∼ 35% of the cell cycle, is associated with more symmetric division site placement. To better characterize the phenotypic determinants of asymmetric cell division, we further used our tracking of the division site backwards in time (Figure 4A) to extract the first observed time *T*_0_ for which the division site gets tracked to mid-cell. The average time at which this occurs corresponded to *T*_0_ at 36% ± 29% of the cell cycle (Table 2), which corresponds to the previously found NETO time at which division is symmetric (Figure 4C). We hence suggest that division site selection occurs on average at this time, and that a relatively later NETO (longer lag phase) results in a more asymmetric division site selection. We also note that as it corresponds to typically 100-120 minutes prior to daughter cell cleavage, and as a potential prediction time for division site selection, this average time for *T*_0_ precedes, or is concomitant with, other morphological and chemical processes known to be involved in division site selection, such as the appearance of the FtsZ ring (∼100 minutes before cleavage) and the formation of the pre-cleavage furrow (∼60 minutes before cleavage), both indicators of septum formation [3].

### 3.5 Mathematical modeling of morphological dynamics

Cell wall architecture influences rod-shaped cell morphology in *M. smegmatis*. Recent evidence demonstrates that expression of L, D-transpeptidases (LDTs) are necessary for maintaining wave-form morphology in *M. smegmatis* [19]. However, it is unclear how mechanistically wave-form morphology is constructed at sub-polar regions or stably maintained near mid-cell. Defining first principles for how a wave-form morphology can be achieved provides critical insights for future molecular screens. Our synthetic approach aims to identify the fewest number of essential factors necessary for creating a wave-form surface morphology.

We propose a minimal mathematical model generating patterns of wave-peaks and wave-troughs. Our model recapitulates the creation of wave-form patterns at sub-polar regions, morphological stability near mid-cell, and wave-form characteristics (amplitude and wavelength) set independently of elongation velocity. Our model is derived from the so-called Schnackenberg model on an 1D evolving domain, a reaction-diffusion process for two “chemical species” (with concentration *u* and *v*) with non linear interactions as Equation (1) :

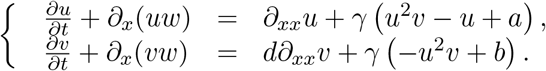

In this model, the rate of change of the concentrations of the chemical species is a mass balance between diffusion of the species (*∂*_*xx*_*u* and *d∂*_*xx*_*v*) and their net production (right terms) with intensity *γ*, with the *∂*_*x*_(*uw*) and *∂*_*x*_(*uw*) accounting for domain deformations. Species *u* and *v* are produced at constant rates *a* and *b*, whereas −*u* is a decaying production term proportional to its concentration. Finally the term *u*^2^*v* represents a non linear autocatalytic reaction between the two species: *u* enhances positively its own production while *v* decays proportionally to the product of its concentration and that of two entities of the species *u*.

Solving the model numerically for a broad range of parameter values (see and SI files Section S2) yielded a wave-form surface morphology reminiscent of our experimental observations of the cell surface of *M. smegmatis*. The wavelength associated with the height patterns varies slightly over time around a fixed value (∼ 0.43), while remaining constant over the cell domain (Figure 5B). Old and new pole wave-form patterns exhibit the same wavelength, independent of the phase of pole elongation dynamics. The wavelength is controlled by the parameter *γ* (Figure S5A, B, E), whereby a decrease in *γ* results in an increase in wavelength. Therefore *γ* can be tuned to reproduce any observed wavelength pattern [20]. We also noted that the diffusion constant *d* controls the sub-polar emergence of wave-form features and the maintenance of stable patterns in the central region. Wave-form patterns disappear at low values of *d* (Figure S5C), whereas fine patterns manifest as *d* increases (Figure S5D). Our simulation displayed slight displacements of the wave-form features over time, which differs from experimental observations. This phenomenon results from the interplay between pattern creation and cell elongation, and could be mitigated by modeling mechanisms of cell wall architecture formation as a function of *u*.

**Figure 5.**
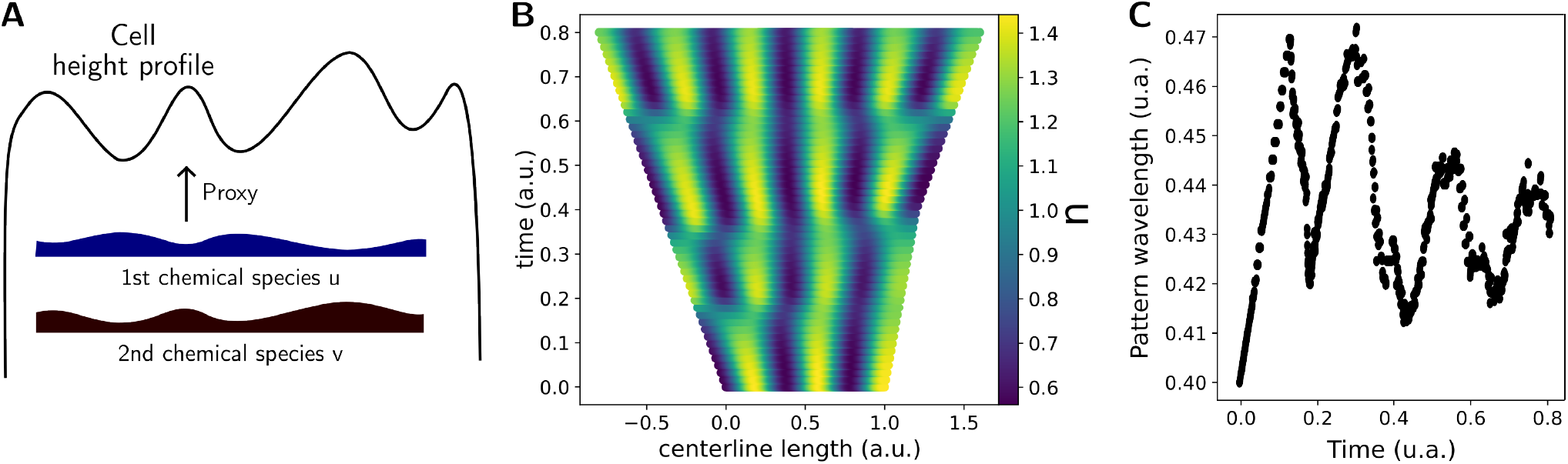
Modeling and simulation of the wave-form surface morphology using a reaction-diffusion system. (A) Representation of the model: hypothetical distributions of both chemical species, *u* and *v*, are depicted along the cell length. Interactions of both chemical species results in wave-form pattern creation. (B) Simulated kymograph of the model representing variable *u* and its patterns. (C) Plot of the simulated pattern wavelength as a function of time.

## 4 Discussion

LTTL-AFM represents an emerging advanced imaging technique, dynamically resolving nanoscale surface morphology of cell surfaces. However, interpretation of LTTL-AFM images has previously been limited by the lack of automated image analysis advanced tools for extracting information about fundamental mycobacterial cell biology from large datasets with minimal human bias [2, 3, 7, 11, 19]. In this paper, we developed and ran a computational pipeline to systematically track and extract novel multi-parametric LTTL-AFM datasets of *M. smegmatis* cells in time-lapse experiments. We used our unique datasets to quantify fundamental biophysical principles governing the establishment and maintenance of *M. smegmatis* morphology. The systematic process of cell segmentation and tracking of cells within LTTL-AFM images reduces human bias and automates the statistical analyses of morphological features.

Rod-shaped (non-sporulating) bacteria are generally described to be composed of one cellular compartment. The extent of sub-cellular compartmentalization can largely be differentiated based on where evolutionarily disparate bacteria harbor growth. Mycobacteria undergo elongation from the poles, much like other actinobacteria and fission yeast [12, 21]. Division occurs near mid-cell. However, emerging evidence provides a refined understanding that the mycobacterial cell surface harbors several functionally distinct zones where discrete fundamental cell processes are achieved [2, 3, 7]. Polar localization of Wag31 is conditionally essential for driving NETO in mycobacteria [2, 22]. In the lag phase, pre-NETO, L,D-transpeptidases are localized to the slow-growing new pole [19]. These studies offer insight into the mechanistic links between biochemical cell processes associated with distinct pole elongation dynamics. The mycobacterial cell surface is equally biophysical in nature. Recent evidence reveals that there exist three mechanically distinct zones at the cell surface [3]. Elongating polar regions harbor mechanically “soft” cell surface material, sub-polar regions of elongating poles harbor zones of hardening cell surface, and mid-cell harbors a zone of biomechanically static and stiff cell surface. Here, we demonstrate that new wave-form morphology is established at sub-polar zones, overlapping with zones where cell surface material is either biomechanically soft or hardening. Future studies using the automated image analysis platform presented here will enable us to determine how mechanistically cell surface biomechanical maturation and molecular determinants control the emergence and maintenance of a wave-form morphology.

Division site selection represents another dynamic and fundamental cell process which occurs at a wave-trough near mid-cell [3]. Previously, it was proposed that wave-troughs represent ‘licensed’ landmarks for division site selection. Wave-troughs are established 1-2 generations prior to division site selection, making them the earliest known morphological features indicating where division occurs in the future [3]. However, it remains unclear how early mycobacteria can mechanistically undertake division site selection. We demonstrate that, by ∼ 36% of the cell cycle, we can infer which wave-trough will harbor division. It is possible that a molecular driver of division site selection occurs within a second trimester of the mycobacterial cell cycle. We additionally demonstrate that NETO timing influences pole growth asymmetry and subsequently the spatial distribution of wave troughs. Previous studies identified *lamA* as a conditionally essential gene in *M. smegmatis* controlling pole growth asymmetry [23]. *LamA* potentially plays a role in controlling the duration of the lag phase, pre-NETO. It is possible that a consequence of *lamA* activity is to influence the spatial distribution of the mycobacterial wave-form morphology and subsequently how early it can be inferred in which wave trough division site selection will occur. Another example of gene associated with asymmetry in cell division is *parB*, with a higher rate of asymmetric cell divisions occurring in *M. smegmatis* Δ*parB* mutant bacilli [3]. Among other mycobacteriae, *M. abscessus* and *M. tuberculosis* exhibit longer cell cycles while maintaining a relatively similar lag phase (pre-NETO) to *M. smegmatis*; these mycobacterial pathogens grow more symmetrically, likely resulting in a differing mean spatial distribution of wave-troughs and difference in how early during the cell cycle a division site selection within a discrete wave-trough can be inferred.

How mechanistically does a wave-form cell surface morphology manifest in *M. smegmatis*? Various biochemical and biophysical parameters, such as the cell wall composition, and turgor pressure, influence morphology in discrete sub-cellular regions [3, 7, 19, 24, 25]. A two species-reaction-diffusion model reveals first principles underlying the dynamic mechanistic process for generating and maintaining a wave-form cell surface morphology. The local concentrations of two distinct physico-chemical species is sufficient to generate an undulating wave-form morphology. How biologically can such a model be encoded? Mechanistically, a crosslinked peptidoglycan layer represents the main load-bearing unit of the cell wall maintaining rod shape by withstanding intracytosolic turgor. Penicillin binding proteins (PBPs), which are responsible for crosslinking nascent peptidoglycan at elongating poles (post-NETO) [26], and their associated cell wall factors may play an important role as their localization corresponds to where cell surface material undergoes mechanical hardening and where wave-form morphology is established [21]. PBP-associated factors may equally play a role in determining locally when a curved cell surface morphology buckles in the opposite direction to produce a wave-form pattern. Utilizing dual LTTL-AFM and optical fluorescence microscopy provides an important opportunity for correlative studies of the relationships between biophysical cell properties and molecular factors.

Here, we provide an image analysis platform that aims to systematically expand the analysis of LTTL-AFM data and provide novel insights for how fundamental cell processes in *M. smegmatis* take place. The automation and expanded ease-of-use of LTTL-AFM offers an expanded potential for the dynamic and multi-parametric visualization of bacterial cell surfaces. The coupling of LTTLAFM derived biomechanical datasets with optical fluorescence microscopy-based visualization of molecular factors provides unique opportunities to bridge discovery-based LTTL-AFM research with more traditional “molecular” and “mechanistic” biology [3]. Defining first principles for cell processes for which molecular mechanisms remain poorly understood remains a critical step towards expanding our understanding of (myco)bacterial cell biology.

## Acknowledgments

K.D.D. and C.S. were supported by a NSERC Discovery grant (RGPIN-2020-05348), NSERC Alliance International grant (585873-2023) and a MITACS PIMS fellowship. H.A.E. was supported by a European Molecular Biology Organization Long Term Fellowship (EMBO LTF 191-2014 and aLTF 750-2016), a Cystic Fibrosis Foundation Pilot and Feasibility Award (#002510I221) at UCSF, a UCSF TB RAP Mentored Scientist Award (R25AI47375), and rapid and standard access proposals with affiliate status at the Molecular Foundry, at Lawrence Berkeley National Laboratory. Work at the Molecular Foundry was supported by the Office of Science, Office of Basic Energy Sciences, of the US Department of Energy under contract no. DE-AC02-05CH11231. A.M. was supported by the Canada Research Chair (Tier 1) in Theoretical and Computational Biology (CRC-2022-00147), the Natural Sciences and Engineering Research Council of Canada (NSERC), Discovery Grants Program (RGPIN-2023-05231), the British Columbia Knowledge Development Fund (BCKDF), Canada Foundation for Innovation – John R. Evans Leaders Fund – Partnerships (CFI-JELF), the British Columbia Foundation for Non-Animal Research, and the UKRI Engineering and Physical Sciences Research Council (EPSRC: EP/J016780/1).

We thank Romain Ageron and Eshan Nirodi for helping coding the wave-features detection and height profiles alignment algorithms, as well as the data structure.

## Competing Interests

HAE is a co-founder of Morphogenesis Bio, Inc. HAE reports no conflicts of interests related to the data or results reported in this paper. The authors declare no competing financial interests.

## Notes

https://osf.io/e86qg/

https://github.com/clementsoubrier/AFM_pipeline_paper_code

### Competing Interest Statement

The authors have declared no competing interest.

## References

[1] Sparks IL, Derbyshire KM, Jacobs Jr WR, Morita YS. Mycobacterium smegmatis: the van-guard of mycobacterial research. Journal of bacteriology 205, e00337–22 (2023).

[2] Hannebelle M, Ven JX, Toniolo C, Eskandarian HA, Vuaridel-Thurre G, McKinney JD, et al. A biphasic growth model for cell pole elongation in mycobacteria. Nature communications 11, 1–10 (2020).

[3] Eskandarian HA, Odermatt PD, Ven JX, Hannebelle M, Nievergelt AP, Dhar N, et al. Division site selection linked to inherited cell surface wave troughs in mycobacteria. Nature microbiology 2, 1–6 (2017).

[4] Eskandarian HA, Nievergelt AP, Fantner GE. Time-Resolved Imaging of Bacterial Surfaces Using Atomic Force Microscopy. Methods Mol Biol 1814, 385–402 (2018).

[5] Binnig G, Quate CF, Gerber C. Atomic Force Microscope. Phys Rev Lett 56, 930–933 (1986).

[6] Voigtländer B. Atomic force microscopy. Springer. (2019).

[7] Eskandarian HA, Chen YX, Toniolo C, Belardinelli JM, Palcekova Z, Hom L, et al. Mechanical morphotype switching as an adaptive response in mycobacteria. Science Advances 10, eadh7957 (2024).

[8] Stringer C, Wang T, Michaelos M, Pachitariu M. Cellpose: a generalist algorithm for cellular segmentation. Nature methods 18, 100–106 (2021).

[9] Ruan X, Murphy RF. Evaluation of methods for generative modeling of cell and nuclear shape. Bioinformatics 35, 2475–2485 (2019).

[10] Cutler KJ, Stringer C, Lo TW, Rappez L, Stroustrup N, Brook Peterson S, et al. Omnipose: a high-precision morphology-independent solution for bacterial cell segmentation. Nature methods 19, 1438–1448 (2022).

[11] Odermatt PD, Hannebelle MT, Eskandarian HA, Nievergelt AP, McKinney JD, Fantner GE. Overlapping and essential roles for molecular and mechanical mechanisms in mycobacterial cell division. Nature physics 16, 57–62 (2020).

[12] Mitchison JM, Nurse P. Growth in cell length in the fission yeast Schizosaccharomyces pombe. Journal of cell science 75, 357–376 (1985).

[13] Schnakenberg J. Simple chemical reaction systems with limit cycle behaviour. Journal of theoretical biology 81, 389–400 (1979).

[14] Barreira R, Elliott CM, Madzvamuse A. The surface finite element method for pattern formation on evolving biological surfaces. Journal of mathematical biology 63, 1095–1119 (2011).

[15] Alnæs M, Blechta J, Hake J, Johansson A, Kehlet B, Logg A, et al. The FEniCS project version 1.5. Archive of numerical software 3 (2015).

[16] Scroggs MW, Baratta IA, Richardson CN, Wells GN. Basix: a runtime finite element basis evaluation library. Journal of Open Source Software 7, 3982 (2022).

[17] Barrata IA, Dean JP, Dokken JS, Habera M, Hale J, Richardson C, et al. DOLFINx: The next generation FEniCS problem solving environment. Zenodo, PREPRINT (2023). DOI: 10.5281/zenodo.10447665.

[18] Unissa AN, Subbian S, Hanna LE, Selvakumar N. Overview on mechanisms of isoniazid action and resistance in Mycobacterium tuberculosis. Infection, Genetics and Evolution 45, 474–492 (2016).

[19] Baranowski C, Welsh MA, Sham LT, Eskandarian HA, Lim HC, Kieser KJ, et al. Maturing Mycobacterium smegmatis peptidoglycan requires non-canonical crosslinks to maintain shape. Elife 7, e37516 (2018).

[20] Madzvamuse A, Thomas RD, Maini PK, Wathen AJ. A numerical approach to the study of spatial pattern formation in the ligaments of arcoid bivalves. Bulletin of mathematical biology 64, 501–530 (2002).

[21] Meniche X, Otten R, Siegrist MS, Baer CE, Murphy KC, Bertozzi CR, et al. Subpolar addition of new cell wall is directed by DivIVA in mycobacteria. Proceedings of the National Academy of Sciences 111, E3243–E3251 (2014).

[22] Santi I, Dhar N, Bousbaine D, Wakamoto Y, McKinney JD. Single-cell dynamics of the chromosome replication and cell division cycles in mycobacteria. Nature communications 4, 2470 (2013).

[23] Rego EH, Audette RE, Rubin EJ. Deletion of a mycobacterial divisome factor collapses single-cell phenotypic heterogeneity. Nature 546, 153–157 (2017).

[24] Palčeková Z, Angala SK, Belardinelli JM, Eskandarian HA, Joe M, Brunton R, et al. Disruption of the SucT acyltransferase in Mycobacterium smegmatis abrogates succinylation of cell envelope polysaccharides. Journal Of Biological Chemistry 294, 10325–10335 (2019).

[25] Hohl M, Remm S, Eskandarian HA, Dal Molin M, Arnold FM, Hürlimann LM, et al. Increased drug permeability of a stiffened mycobacterial outer membrane in cells lacking MFS transporter Rv1410 and lipoprotein LprG. Molecular microbiology 111, 1263–1282 (2019).

[26] Vollmer W, Blanot D, De Pedro MA. Peptidoglycan structure and architecture. FEMS microbiology reviews 32, 149–167 (2008).

