## Supporting Material for "Quantifying the organization and dynamics of *M. smegmatis* morphology from Long-Term Time-Lapse Atomic Force Microscopy"

The Supplemental Information contains:

- S1 Supplementary material detailing data processing pipeline (Protocol)
- S2 Supplementary numerical scheme for the implementation of the Schnackenberg model on evolving domains
- S3 Supplementary Figures S1–S5
- S4 Supplementary table of the datasets used in the figures

### S1 Protocol

In this section, we describe the image analysis pipeline implemented to extract single cell morphological trajectories from LLTL-AFM raw data. The microscope data is first extracted and filtered, then segmented to detect cells which are subsequently tracked. Geometrical features are then extracted and height profiles are created. Finally, height profiles are aligned and peaks and troughs are detected and tracked.

#### S1.1 Image processing and single cell segmentation

Microscopes log files contain several channels, each representing a physical property such as height, stiffness, peak force error and so forth. Images, as well as key parameters, such as units, image rotation angle or time, are extracted from the log files using the pySPM python package. For each dataset, images are first re-sized into common dimensions and resolution. Image scars are then detected and corrected. In order to compensate the drift caused by the hysteresis of the AFM piezo-electric sensor [1], backward and forward images are aligned using an intensity based rigid body method [2].

Segmentation of each frame is performed by using the python package Cellpose [3] with optimized hyper parameters. To reinforce contrast and thus improve the segmentation, backward and forward images of the peak force error channel are superimposed to the height channel. Once segmented, the height channel is flattened using the 2D linear interpolation of the background image. Masks resulting in undersized segmentation or presenting large saturation areas are filtered out.

#### S1.2 Lineage tracing

We developed and implemented a tracking algorithm to extract lineage information and discard high defect data. To do so, a tracking algorithm has been developed under the assumption of small variations between the frames. In particular, if the segmented masks are not impacted by an artifact or defect, the size, geometry and cell position are similar between two frames. Since the rotation angle between two frames is known, a translation vector is first computed. Since noise and major defects in the height data might impact intensity based alignment, we first binarize the images (0 if the pixel is outside of a mask and 1 otherwise) and then align using intensity based methods.

Once the translation vector is computed, every mask within a certain time window is used to build an asymmetrical score matrix. The score $A_{i,j}$ (a float between 0 and 1) between two masks $m_i$ and $m_j$ is computed as the intersection area of the 2 masks relative to the first mask area:

$$A_{i,j} = \frac{\text{area}(m_i \cap m_j)}{\text{area}(m_i)}.$$

From the score matrix, a link oriented graph is produced, from mother to daughter. If the shape and position of the mask does not change much between two images, the corresponding scores $A_{j,i}$ and $A_{i,j}$ are high. Masks $i$ and $j$ with corresponding scores above a threshold $t_l$

are linked, i.e. if $A_{i,j} \geq t_l$ and $A_{j,i} \geq t_l$. Division of a mother cell $m_i$ into two daughter cells of the cell frame $m_j, m_k$ is also detected when the following holds

$$A_{k,i} \geq t_l \text{ and } A_{j,i} \geq t_l \text{ and } A_{i,k} + A_{i,j} \geq t_l.$$

In other words, a link is created between the mother and daughter if the union of the daughters and the mother have both similar shapes. Note that the created graph is not a tree in general. Roots, branching and terminal points are then detected. Finally, lineages (representing pictures of a cell from birth to division or death) are constructed as the deepest graph going from a root or a branching point. These graphs are filtered based on their size and quality and re-glued together at division point, making sure that two daughters have non intersecting masks.

#### S1.3 Centerlines extraction and alignment

An important geometrical feature of *M. smegmatis* cells is their height profile, which represent the height along the centerline of the cell. Centerlines are computed from the skeletonization of the segmented masks using the scikit-image python library [4]. Due to the roughness and geometry of the masks, computed skeletons stops far from the cell boundary and may have several branches. They are then pruned and extended. Skeleton with many branches usually indicates a problem in the segmented masks and are used to discard them. Height profile are then derived using centerlines as curvilinear abscissa. In order to produce the kymographs, height profiles from a same lineage are then aligned. Suppose that profiles $f$ and $g$ are defined on the intervals $I_f, I_g$ respectively. We suppose that $I_g$ is smaller then $I_f$. We define $\delta(I_f, I_g)$ as the length of the interval $I_g \backslash I_f$, that is to say the length of $I_g$ that is not in $I_f$. We align the height profiles optimizing:

$$\min_{\substack{h \in \mathbb{R} \\ I_g, \delta(I_f, I_g) \leq l_0}} \text{quantile}_r \left\{ \frac{\int_I (f-g-h)^2}{\int_I dx}, \quad I \text{ interval in } I_f \cap I_g \text{ of lenght } l_1 \right\} + \epsilon \delta(I_f, I_g),$$

where $\text{quantile}_r$ compute the $r$ quantile of a distribution, $r, \epsilon, l_0, l_1$ are fixed parameters. If the parameters are well chosen, this metric will align height profiles focusing on region with similar geometry and discarding regions where the geometry highly evolve (typically pole regions that are growing).

#### S1.4 Features detection and tracking

Each height profile present feature (peaks and troughs) which are more significant than the characteristic noise. After Gaussian smoothing of the height profiles, local extrema are detected, where the derivative of the curve changes sign. Features are then filtered depending on their amplitude and distance between each other, and characterized as peaks and troughs. Inside a lineage, features are tracked using their position on the aligned height profiles. Peaks and troughs are tracked separately based on nearest neighbors over several frames. More precisely, partial lineages are first linking point $x$ and $y$ such that $x$ is the nearest neighbor of $y$ and $y$ is the nearest neighbor of $x$. Partial lineages are then assembled and linked to free features such that peaks and troughs lineages do not intersect.

### S2 Implementation of the reaction-diffusion system

In this section, we describe the numerical scheme used to simulate pattern formation on the cell. We assume that the domain deforms following a bi-phasic and asymmetrical dynamics, such that the old pole grows at a constant rate, and the new pole growth rate is first at low value, and then switches to a higher value. We can thus compute the material velocity vector field $\mathbf{w}$ of the domain $\Omega$, that is to say the point $\mathbf{x}$ at time $t$ has the velocity $\mathbf{w}(\mathbf{x}, t)$. Let us discretize in time and space. We now discretize in time using finite differences of step size $\Delta t$ and in space using a mesh size $h$. Let $\Omega_h^n$ the discretized domain at time $n\Delta t$ and $(\chi_i^n)_i$ a finite element basis that follows the material velocity:

$$u(\cdot, n\Delta t) = \sum_i U_i^n \chi_i^n,\ v(\cdot, n\Delta t) = \sum_i V_i^n \chi_i^n.$$

The numerical scheme to solve (1) on the evolving domain becomes [5, 6]:

$$\sum_i \int_{\Omega_h^{n+1}} \left( \frac{1}{\Delta t} U_i^{n+1} - \gamma (U_i^{n+1})^2 V_i^{n+1} - \gamma U_i^{n+1} - \gamma a \right) \chi_i^{n+1} \chi_j^{n+1}$$
$$+ \int_{\Omega_h^{n+1}} U_i^{n+1} \nabla_{\Omega_h^{n+1}} \chi_i^{n+1} \nabla_{\Omega_h^{n+1}} \chi_j^{n+1} = \sum_i \int_{\Omega_h^n} \frac{1}{\Delta t} U_i^n \chi_i^n \chi_j^n,$$

and

$$\sum_i \int_{\Omega_h^{n+1}} \left( \frac{1}{\Delta t} V_i^{n+1} + \gamma (U_i^{n+1})^2 V_i^{n+1} - \gamma b \right) \chi_i^{n+1} \chi_j^{n+1}$$
$$+ \int_{\Omega_h^{n+1}} d V_i^{n+1} \nabla_{\Omega_h^{n+1}} \chi_i^{n+1} \nabla_{\Omega_h^{n+1}} \chi_j^{n+1} = \sum_i \int_{\Omega_h^n} \frac{1}{\Delta t} V_i^n \chi_i^n \chi_j^n.$$

This system was implemented in finite elements software which does not enable two different meshes to be used ( for example $\Omega_h^{n+1}$ and $\Omega_h^n$ ). However we can extract at any time the mass matrix

$$M_{i,j}^n = \int_{\Omega_h^n} \chi_i^n \chi_j^n,$$

which is always symmetric and invertible (positive definite). Thus we have:

$$\begin{aligned} \sum_i \int_{\Omega_h^n} \frac{1}{\Delta t} U_i^n \chi_i^n \chi_j^n &= \frac{1}{\Delta t} (M^n U^n)_j \\ &= \frac{1}{\Delta t} (M^{n+1} (M^{n+1})^{-1} M^n U^n)_j \\ &= \sum_i \int_{\Omega_h^{n+1}} \frac{1}{\Delta t} ((M^{n+1})^{-1} M^n U^n)_i \chi_i^{n+1} \chi_j^{n+1}, \end{aligned}$$

and the final numerical scheme becomes:

$$
\begin{aligned}
&\sum_i \int_{\Omega_h^{n+1}} \left(\frac{1}{\Delta t}U_i^{n+1} - \frac{1}{\Delta t}((M^{n+1})^{-1}M^nU^n)_i\right)\chi_i^{n+1}\chi_j^{n+1} \\
&-\sum_i \int_{\Omega_h^{n+1}} \gamma\left((U_i^{n+1})^2V_i^{n+1} + U_i^{n+1} + a\right)\chi_i^{n+1}\chi_j^{n+1} \\
&= -\sum_i \int_{\Omega_h^{n+1}} U_i^{n+1}\nabla_{\Omega_h^{n+1}}\chi_i^{n+1}\nabla_{\Omega_h^{n+1}}\chi_j^{n+1}, \\
&\text{and} \\
&\sum_i \int_{\Omega_h^{n+1}} \left(\frac{1}{\Delta t}V_i^{n+1} - \frac{1}{\Delta t}((M^{n+1})^{-1}M^nV^n)_i + \gamma((U_i^{n+1})^2V_i^{n+1} - b)\right)\chi_i^{n+1}\chi_j^{n+1} \\
&= -\sum_i \int_{\Omega_h^{n+1}} dV_i^{n+1}\nabla_{\Omega_h^{n+1}}\chi_i^{n+1}\nabla_{\Omega_h^{n+1}}\chi_j^{n+1}.
\end{aligned}
\tag{S1}
$$

Note that this system is non linear because of the term $(U_i^{n+1})^2V_i^{n+1}$. Thus a non linear optimization step is needed to solve Eq.(S1). In practice, a simple Newton scheme converges quickly enough to be used [7, 8]. We can also approximate the non linear part with the previous value of the function:

$$(U_i^{n+1})^2V_i^{n+1} \approx U_i^{n+1}U_i^nV_i^n \approx (U_i^n)^2V_i^{n+1}.$$

This gives us the linear IMEX (implicit-explicit) scheme:

$$
\begin{aligned}
&\sum_i \int_{\Omega_h^{n+1}} \left(\frac{1}{\Delta t}U_i^{n+1} - \frac{1}{\Delta t}((M^{n+1})^{-1}M^nU^n)_i\right)\chi_i^{n+1}\chi_j^{n+1} \\
&-\sum_i \int_{\Omega_h^{n+1}} \gamma\left(U_i^{n+1}U_i^nV_i^n + U_i^{n+1} + a\right)\chi_i^{n+1}\chi_j^{n+1} \\
&= -\sum_i \int_{\Omega_h^{n+1}} U_i^{n+1}\nabla_{\Omega_h^{n+1}}\chi_i^{n+1}\nabla_{\Omega_h^{n+1}}\chi_j^{n+1}, \\
&\text{and} \\
&\sum_i \int_{\Omega_h^{n+1}} \left(\frac{1}{\Delta t}V_i^{n+1} - \frac{1}{\Delta t}((M^{n+1})^{-1}M^nV^n)_i + \gamma((U_i^n)^2V_i^{n+1} - b)\right)\chi_i^{n+1}\chi_j^{n+1} \\
&= -\sum_i \int_{\Omega_h^{n+1}} dV_i^{n+1}\nabla_{\Omega_h^{n+1}}\chi_i^{n+1}\nabla_{\Omega_h^{n+1}}\chi_j^{n+1}.
\end{aligned}
$$

We analyzed the simulation results for model parameters in the following ranges: $a \in [0.1, 5]$, $b \in [0.1, 5]$, $d \in [10, 100]$ and $\gamma \in [10, 1000]$. We extracted values where patterns appear with dynamics agreeing with our experimental results. If not otherwise stated, parameter used in Figure 5 and S5 are $a = 0.1$, $b = 0.9$, $d = 10$ and $\gamma = 800$.

### S3 Supplementary Figures

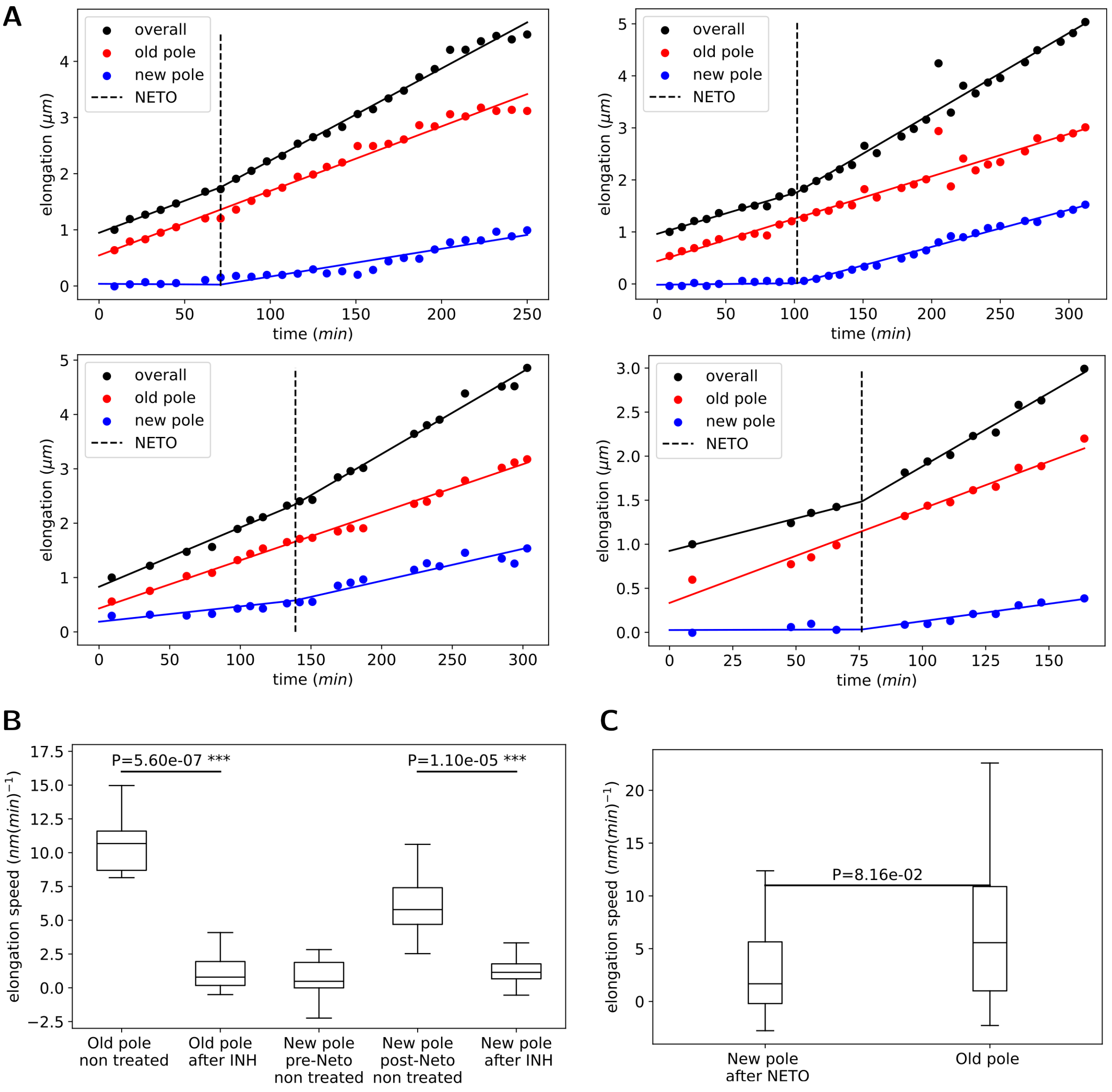


**Figure S1.** Bi-phasic polar elongation in *M. smegmatis*. (A) Graph representing elongation trajectories for each pole and the overall cell, as well as the NETO. (B) Polar elongation speed comparison for wild type and INH exposed bacteria. (C): Comparison of polar elongation between old pole and new pole after NETO.

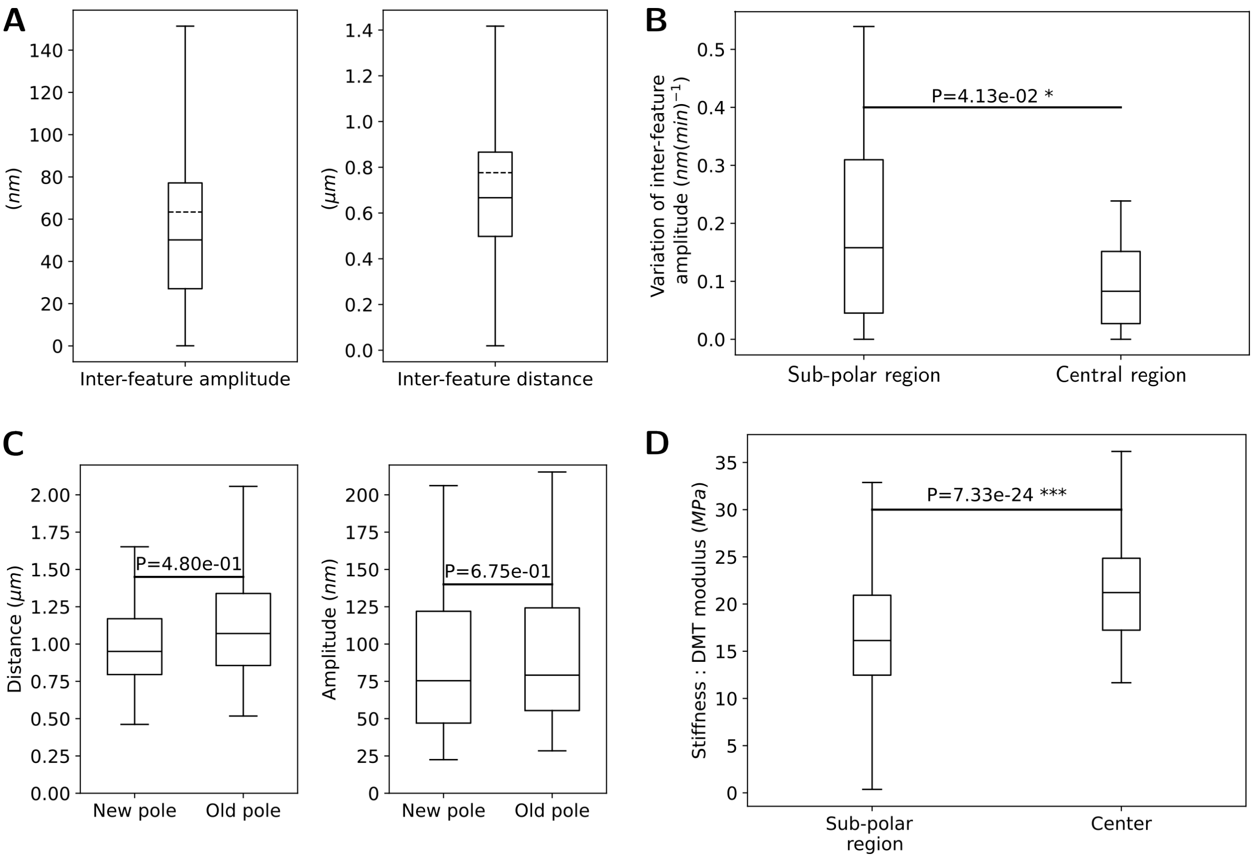


**Figure S2.** Morphological pattern properties in different cell regions. (A) Pattern inter-features distance and amplitude. (B) Comparison of inter-feature amplitude variation between sub-polar and central cell regions. (C) Pattern distance and amplitude at the old and new pole. (D) Comparison of cell stiffness between sub-polar and central cell regions.

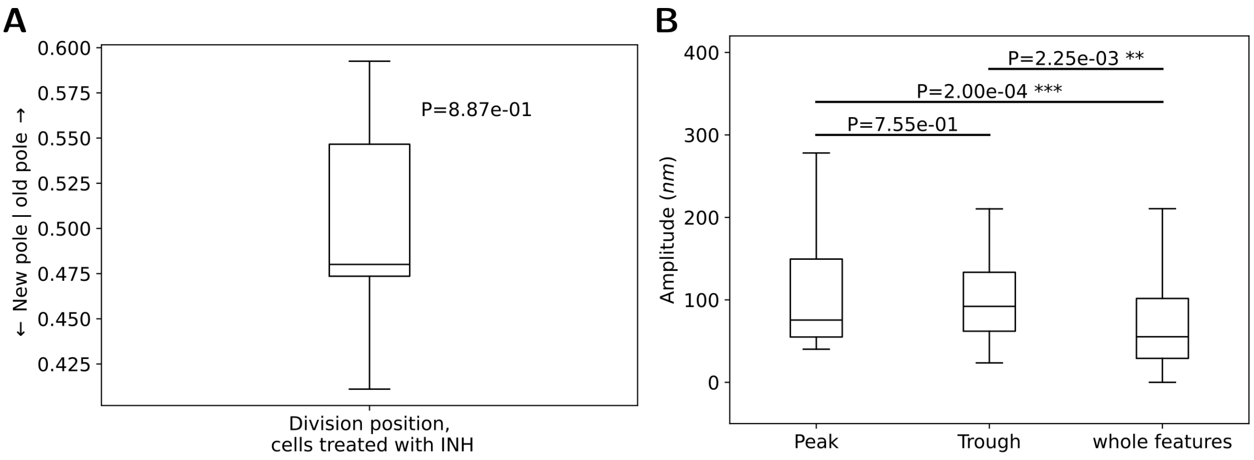


**Figure S3.** Characterization of division site position and their closest features. (A) Division site position of INH treated cells. (B) Amplitude of nearest peaks and troughs to division site, compared to overall features amplitude.

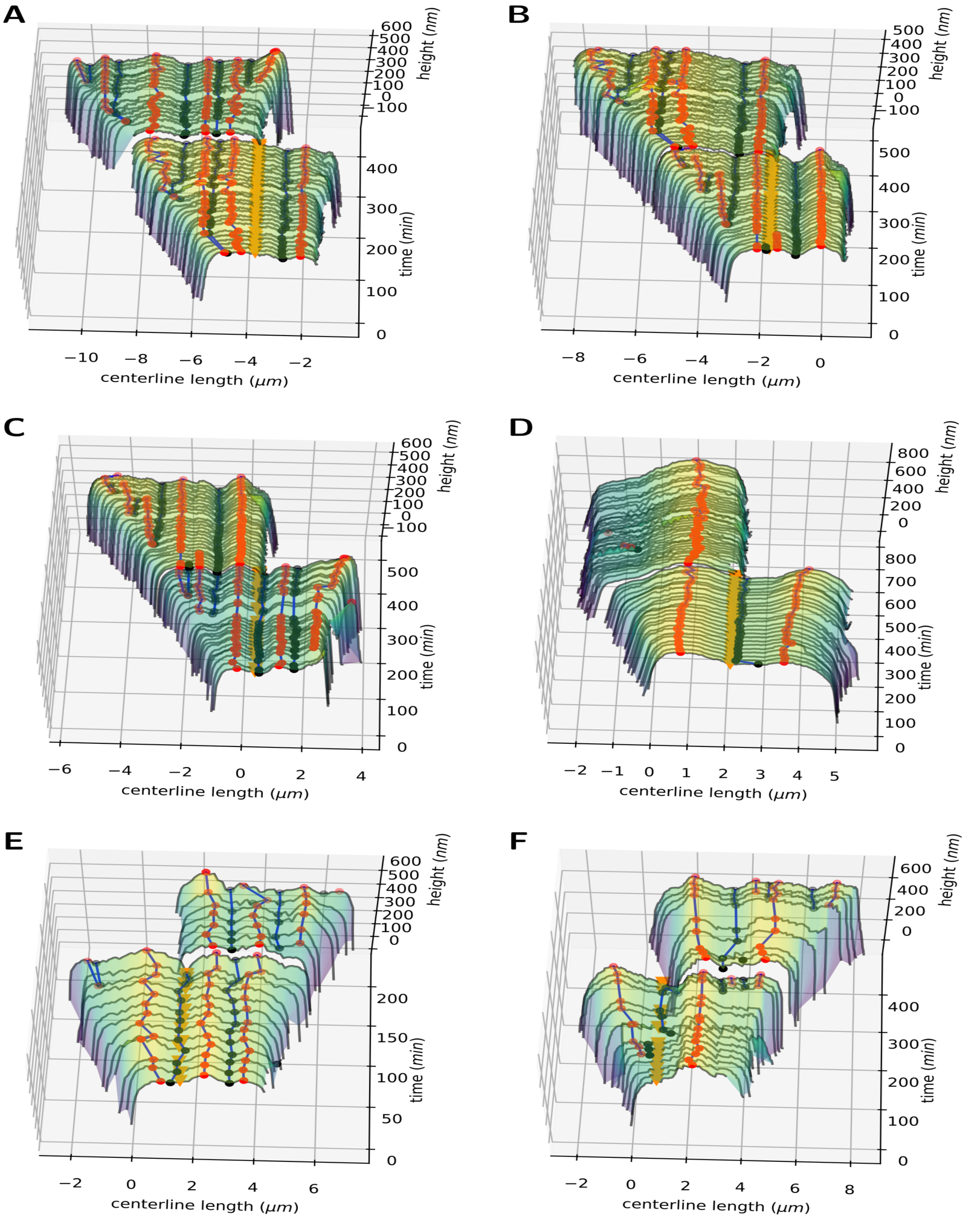


**Figure S4.** Examples of kymographs produced by our pipeline. (A),(B), (C) and (E) Kymograph of wild types untreated cells. (D) Kymograph of a Δ*ltd6* mutant that develops swelling. (F) Kymograph of wild type cells treated with Ciprofloxacin.

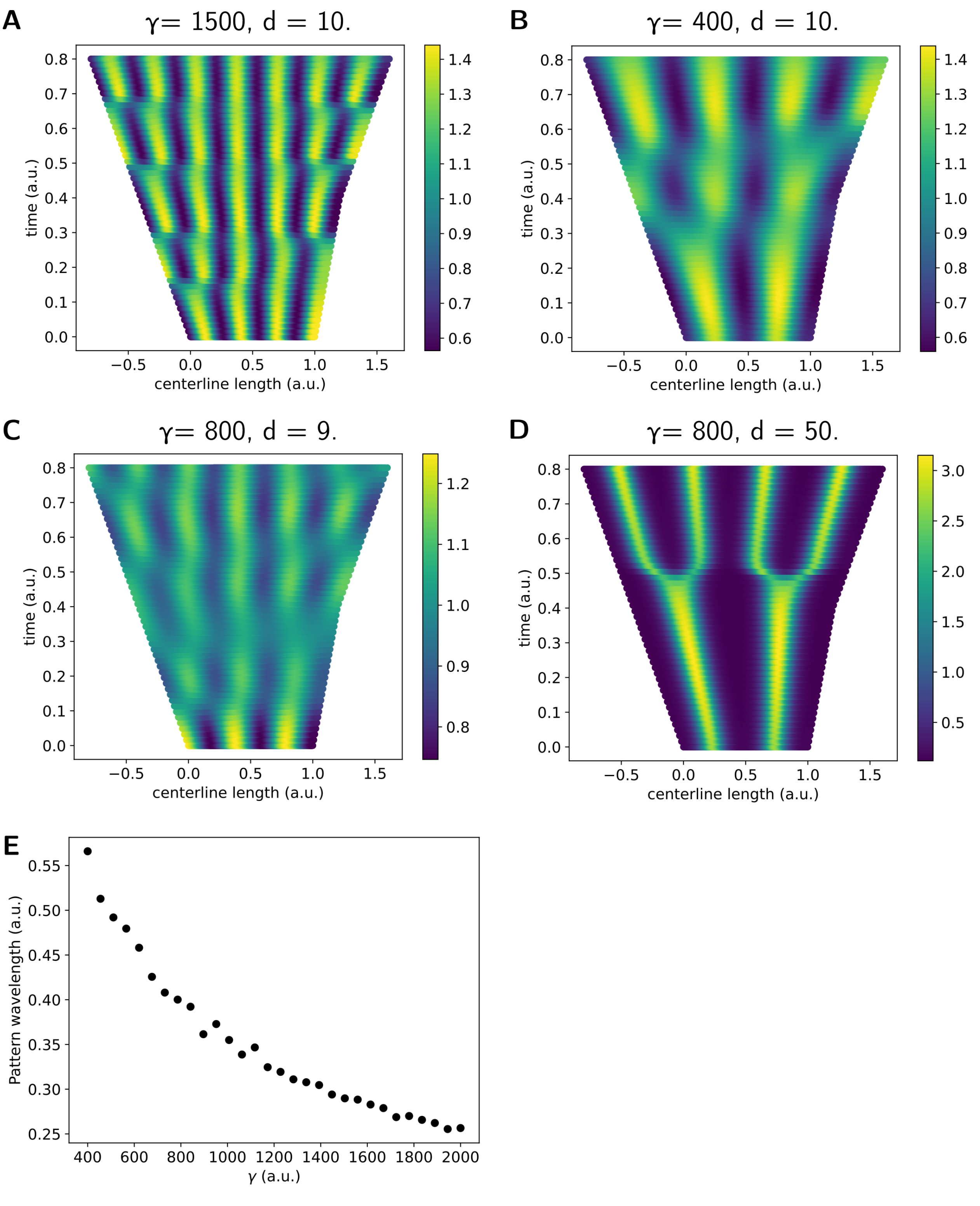


**Figure S5.** Parameter analysis for pattern formation and simulations of our reaction-diffusion system. (A), (B), (C), (D) Simulation of the reaction-diffusion system with different parameters $\gamma$ and $d$. (E) Graph of pattern wavelength as a function of parameter $\gamma$.

### S4 Supplementary Table

| Dataset | Figure / Table | Frames,<br>Detected masks,<br>Tracked cell |
|---|---|---|
| WT best | F.2, F.3, F.4 (B),<br>T.1, F.S1 (A, B), F.S2 (A, B, D)<br>F.S3 (B), F.S4 (A, B, C) | 135, 423, 16 |
| WT un-treated | T.2, F.4 (C), F.S1 (C), F.S2 (C) | 402, 12526, 336 |
| INH treated | F.S1 (B), F.S3 (A), F.S4 (D) | 89, 3825, 86 |

**Table S1.** Datasets used for the statistical analysis. The data we used was previously published in [9, 10]. “WT best” dataset contains high quality images of untreated wild type bacteria, with a low number of cells. “WT untreated” includes “WT best” and other untreated wild types datasets of lower quality, but with a higher number of individual cells. It is used to statistically confirm trends that were detected in “WT untreated” . “INH treated” contains wild type cells that have been treated with isoniazid.

### REFERENCES


1. Rodriguez-Fortun JM, Orus J, Alfonso J, Buil F, Castellanos JA. Hysteresis in Piezoelectric Actuators: Modeling and Compensation. IFAC Proceedings Volumes **44**, 5237–5242 (2011). 18th IFAC World Congress.

2. Brown LG. A survey of image registration techniques. ACM computing surveys (CSUR) **24**, 325–376 (1992).

3. Stringer C, Wang T, Michaelos M, Pachitariu M. Cellpose: a generalist algorithm for cellular segmentation. Nature methods **18**, 100–106 (2021).

4. van der Walt S, Schönberger JL, Nunez-Iglesias J, Boulogne F, Warner JD, Yager N, et al. scikit-image: image processing in Python. PeerJ **2**, e453 (2014).

5. Dziuk G, Elliott CM. Finite elements on evolving surfaces. IMA journal of numerical analysis **27**, 262–292 (2007).

6. Barreira R, Elliott CM, Madzvamuse A. The surface finite element method for pattern formation on evolving biological surfaces. Journal of mathematical biology **63**, 1095–1119 (2011).

7. Lakkis O, Madzvamuse A, Venkataraman C. Implicit–explicit timestepping with finite element approximation of reaction–diffusion systems on evolving domains. SIAM Journal on Numerical Analysis **51**, 2309–2330 (2013).

8. Madzvamuse A, Chung AH. Fully implicit time-stepping schemes and non-linear solvers for systems of reaction–diffusion equations. Applied Mathematics and Computation **244**, 361–374 (2014).

9. Eskandarian HA, Odermatt PD, Ven JX, Hannebelle M, Nievergelt AP, Dhar N, et al. Division site selection linked to inherited cell surface wave troughs in mycobacteria. Nature microbiology **2**, 1–6 (2017).

10. Odermatt PD, Hannebelle MT, Eskandarian HA, Nievergelt AP, McKinney JD, Fantner GE. Overlapping and essential roles for molecular and mechanical mechanisms in mycobacterial cell division. Nature physics **16**, 57–62 (2020).
